# Incidence and Antibiotic Susceptibility Profile of *Plesiomonas shigelloides* Isolated from Fish, Fish Storage water and Fish seller’s towel

**DOI:** 10.1101/2024.05.01.592048

**Authors:** Temitope Deborah Agboola, Monica Oluwatoyin Oguntimehin, Isaac Olaoluwa Obadofin, Oluwapelumi Ruth Oyeneye

**Affiliations:** Department of Biological Sciences (Microbiology Program), Olusegun Agagu University of Science and Technology, Okitipupa, Ondo State Nigeria

**Author notes:** **Corresponding Author:** Temitope Deborah AGBOOLA; /+2348056061545.

**Keywords:** *Plesiomonas shigelloides*, *Clarias gariepinus*, Towel, Antibiotics Susceptibility, Resistant phenotypes

## Abstract

This study evaluated the incidence of *Plesiomonas shigelloides* and their antibiogram pattern in fish storage water, fish and fish seller’s towel swab collected from two major markets in Southern part of Ondo State, Nigeria. About 158 presumptive *Plesiomonas shigelloides* were recovered from the samples collected from the two markets and only 58 (31.6%) were positive using PCR (polymerase chain reaction) method. Using the disc diffusion method, confirmed isolates were assessed for their antibiogram profiles against 15 antibiotics and multiple antibiotic resistance phenotypes. Resistance of the isolates against the antibiotics followed the order: Erythromycin (85%), Ampicillin (83%), Ceftazidime (81%), Cefuroxime (71%), Tetracycline (67%), Meropenem and Vancomycin (45%), Amoxicillin and Streptomycin (43%), Trimethoprim (41%), Ciprofloxacin and Chloramphenicol (34%), Gentamicin (26%), Cotrimoxazole (22%) and Amikacin (21%). Conversely, all the isolates did not show 100 % susceptibility to any antibiotics, and susceptibility against the antibiotics follows the order: Amikacin (79%), Cotrimoxazole (78%), Gentamicin (74%), Ciprofloxacin and Chloramphenicol (66%), Trimethoprim (59%), Amoxicillin and Streptomycin (57%), Meropenem and Vancomycin (55%), Tetracycline (33%), Cefuroxime (29%), and exhibited less than 20% susceptibility to other antibiotics. The multiple antibiotic-resistance indices of the organism were higher than the accepted threshold of 0.2. This is the first report on assessing the ease at which *P. shigelloides* in *Clarias gariepinus* sold in the market can be transmitted. We concluded that this pathogen with multiple antimicrobial resistant phenotypes can be easily transmitted consequently, a public health threat meanwhile, Amikacin, Cotrimoxazole and Gentamicin are important antibiotics that could be used against the pathogen.

## INTRODUCTION

*Plesiomonas shigelloides* is a gram-negative, facultatively anaerobic rod that does not produce spores and has been linked to gastroenteritis epidemics that are transmitted by food and water. *P. shigelloides*, a type of bacteria found in aquatic environments, has reportedly been linked to severe mortalities in both wild and cultivated fish populations worldwide (Wang *et al*., 2020). A food with excellent nutritional value and a necessary component of the human diet, catfish (*Clarias gariepinus*) is widely produced around the world, including Nigeria (Zhang *et al*., 2019; Fakorede *et al*., 2020). With the rapid advancement of farming methods in Nigeria, catfish has emerged as one of the most significant freshwater fish species for commerce and has generated substantial profits in recent years (Fakorede *et al*., 2020). Fish skin, gills, and intestines frequently harbor microorganisms like *P. shigelloides*, which typically degrade fish quality. According to some sources, *P. shigelloides* is the only species in the genus (Habs and Schubert, 1962) and is a part of the growing group of emerging and major foodborne and waterborne pathogens that have been found in a variety of sources, such as water, crabs, fish, and oysters (Aldova, 1999; Nimmervoll *et al*., 2011; Adesiyan *et al*., 2019). It was projected that *P*. *shigelloides* and other pathogens would be present in water bodies because of their proximity to human settlement and as a result of anthropogenic activity (Agboola *et al*., 2021; Fuentes-Valencia *et al*. 2022). The organism has been assigned to the family of which it is the only oxidase-positive member because molecular investigations have demonstrated that it has a tight relationship with the *Enterobacteriaceae* family. Aquatic environments and the animals that live there have been linked to this bacterium most often (Gonzalez-Rey *et al*., 2003; Alexander *et al*., 2016). *Plesiomonas shigelloides* is recognized as a common cause of human enteric infections and has gained importance because it typically results in cholera-like diarrhea in infected individuals, especially after consuming tainted water and seafood (Schneider *et al*., 2009; Bodhidatta *et al*., 2010; Alexander *et al*., 2016). This bacterium also causes a number of extra intestinal conditions that are highly lethal, including pneumonia, septicemia, and meningitis, particularly in people with impaired immune systems (Ozdemir *et al*., 2010; Klatte *et al*., 2012; Miyoshi, 2013). Also, the pathogen has been connected to outbreaks and sporadic episodes of diarrhea worldwide (Bodhidatta *et al*., 2010; Bowman *et al*., 2016). According to Nwokocha and Onyemelukwe (2014), *P. shigelloides* is a gastrointestinal pathogen and may cause diarrhea in Nigeria more frequently than is currently thought. *Plesiomonas shigelloides* infections may be misdiagnosed in the lab as being caused by other prevalent diarrheagenic microbes because there is little information available on the diarrhea associated with this organism in Nigeria. Geographically, the prevalence of gastroenteritis caused by *P. shigelloides* has been observed to vary, with Africa and Southeast Asia having the greatest rates. Seasonal fluctuations, poor hygiene habits, and a lack of portable water sources have been blamed for this (Cortes-Sanchez *et al*., 2021). Extraintestinal infections such as acute cholecystitis, cellulitis, endophthalmitis, septicemia, meningitis, osteomyelitis, septic arthritis and spontaneous bacterial peritonitis are less prevalent and mostly affect patients in immunoincompetent or immunocompromised status (Adesiyan *et al*., 2019). *Plesiomonas shigelloides* is a pathogen that most usually affects fish, where it results in hemorrhage around the vent, protruding anuses, and inappetence (Paul *et al*., 1990). According to Rajagopalan *et al*. (2014), Cichlid fish infected with *P. shigelloides* died at a rate of 100%. Freshwater *P. shigelloides* is a well-known common bacteria, and numerous countries have reported studies of this bacteria’s resistance to antibiotics including aminoglycosides and penicillin (Avison *et al*., 2000; Adesiyan *et al*., 2019) while numerous studies have also found evidence of resistance to drugs including erythromycin, streptomycin, sulphamethoxazole, and tetracycline. (Wong *et al*., 2000; González-Rey *et al*., 2004). Little or no investigation has documented the presence of *P. shigelloides* in aquatic animal as well as the hand towel of fresh fish sellers in Southwestern Nigeria to the best of our knowledge and we present the incidence and antibiogram pattern *P. shigelloides* in fresh fish commonly sold in this area for the first time and the incidence of the pathogen in fish seller’s towel swab in order to offer helpful information on the ease of transmission of this emerging pathogen and how to prevent its associated infection.

## Materials and Methods

### Description of Study Sites

This study was conducted in Okitipupa and Igbokoda areas in the Southern zone of Ondo State, Southwest, Nigeria. The selected study areas are the most common sources of fresh fish for the community’s needs. One of the six States of Southwest Nigeria is Ondo State. The States of Ekiti and Kogi border the State on its northern and western borders, respectively. Additionally, Ondo State is bordered by the Delta and Edo State in the east and by the Atlantic Ocean in the south (Thompson and Mafimisebi, 2014). The State is said to comprise 18 Local Government Areas (LGA) having about 3.4 million inhabitants (NPC, 2007). Ondo State had been reported to have three (Noorlis *et al*., 2011) distinct zones ecologically; to the south is the mangrove forest, the rain forest lies at the center and to the north is guinea savannah. The coordinate of the Okitipupa and Igbokoda sampled sites are 6° 29’ 58^’’^ N, 4° 47^’^ 12^’’^E and 6° 21’ 19’’N, 4° 47’ 53’’ respectively.

### Sampling and isolation of presumptive *P. shigelloides*

A total of twelve (12) fish storage water effluents were collected aseptically from fish sellers using sterile 1000 mL plastic bottles, 12 fish samples were purchased, kept inside zip-lock bags and 10 fish seller’s towels each from Okitipupa and Igbokoda areas of Ondo State. Samples were transferred using ice to the Microbiology laboratory at Olusegun Agagu University of Science and Technology for analysis within six hours of collection. The samples were processed in accordance with the American Public Health Association’s guidelines (APHA, 2005). Fish storage water samples, and fish seller’s towel swab samples and fish homogenates were serially diluted after being enriched in alkaline peptone water (pH 8.6) and incubated at 37 °C for 24 h. Following dilution, 0.1 ml of each sample’s dilutions 3 and 4 were plated on clearly labeled dry plates of Inositol Brilliant green bile salt agar (Conda Pronadisa, Spain). The samples were immediately spread uniformly across the surface of the agar plate using a sterilized glass spreader. The plates were then incubated for 24 to 48 h at 37 °C while being inverted. On each plate, 5 – 10 pinkish colonies were chosen as potential *P. shigelloides* colonies. Only Gram-negative, oxidase positive isolates were chosen after been purified on nutrient agar plate and kept on nutrient agar slants for further analysis.

### DNA Extraction

The boiling procedure was used for DNA extraction (Jeong *et al*., 2015; Agboola *et al*., 2022). On non-selective agar plates, presumptive *P. shigelloides* colonies were subcultured and incubated for 18 to 24 h at 37 °C. Distinct colonies of pure culture were picked into 100 μl sterilized distilled water, vortex and boiled at 100 °C for 15 min and later centrifuged at 13,000 rpm for 15 min. The resulting supernatants were stored at −20 °C in Eppendorf tube as template DNA for PCR analysis.

### Confirmation of Plesiomonas shigelloides isolates

By using the primer sets PS23FW3/PS23RV3 created by González-Rey *et al*. (2000), which amplifies a 284 bp sequence of the 23SrRNA gene (PS23FW3: 5′-CTCCGAATACCGTAGTGCTATCC-3) and (PS23RV3: 5′-CTCCCCTAGCCCAATAACACCTAAA-3), and the polymerase chain reaction technique, the probable *P. shigelloides* identification was validated.

With a few adjustments, the PCR conditions utilized were those stated by Adesiyan *et al*. (2019). About 12.5 μl of PCR master mix (Inqaba Biotech, SA), 6.5 μl of nuclease-free water, 1 μl of each oligonucleotide primers, and 5 μl of DNA template made up the 25 μl total reaction volume. The PCR reaction’s protocols called for an initial denaturation step of 5 min at 95 °C, followed by denaturation, annealing, and extension phases of 1 min each at 94 °C, 68 °C, and 72 °C, for 2 min respectively, and a final extension step of 10 min at 72 °C after 35 cycles. The amplicon’s 5 μl were loaded on a 1.5% agarose gel stained with ethidium-bromide dye to perform the electrophoresis. PCR results were observed using gel documentation tools (UVsave, Applied Bioscience, UK). For molecular size calibration on the gel, a DNA ladder of 100 bp was utilized, and the electrophoresis ran at 100 V for 45 minutes. As a benchmark, *P. shigelloides* DSMZ 8224 was employed. Six out of the confirmed isolates were sequenced for further confirmation.

### Antibiotic Susceptibility Testing

The disc diffusion method was used to perform an antibiotic susceptibility test (Kirby *et al*., 1966). From an 18-hour-old culture of the test organism, distinct colonies were selected and put in tubes with 5 ml of 0.85% physiological saline. The resulting mixture’s turbidity was measured against 0.5 McFarland (equivalent to 1.5 × 10^8^ CFU) reference solutions. Mueller Hinton agar plates were covered with 0.1 ml of the standardized suspension using a sterile swab, and they were allowed to dry before an antibiotic disc was placed on top of them. Fifteen antibiotics (Mast Diagnostics, UK) were chosen for the test and the antibiotics include Tetracycline (30µg), Cotrimoxazole (25µg), Amikacin (30µg), Ampicillin (10 µg), Streptomycin (30μg), Gentamicin (10µg), Vancomycin (30µg); Cefuroxime (30µg), Ceftazidime (30µg), Trimethoprim (5μg), Amoxicillin (25μg), Chloramphenicol (30µg), Meropenem (10µg), Erythromycin and Ciprofloxacin (5µg). On Mueller Hinton agar plates, antibiotic discs were put, and they were then incubated at 37 °C inverted for 24 hours. According to the Clinical and Laboratory Standards Institute’s (CLSI, 2018) advice for interpreting the zone diameter, the zones of inhibition were determined and recorded as susceptible (S), resistant (R), or intermediate (I). The results of the antibiotic susceptibility testing were used to determine the frequency, proportion, and pattern of antibiotic resistance. For isolates that displayed resistance to more than two antibiotics at each sample site, multiple antibiotic-resistant phenotypes (MARPs) were also calculated.

According to Titilawo *et al*. (2015), the MAR index is a crucial indicator for locating contamination that may be hazardous to humans. Its mathematical expression is as follows:

> MARindex = a/b
>
> a = number of antibiotics to which isolates were resistant.
>
> b = total antibiotics to which an individual isolate was tested.

## RESULTS

### PCR Confirmation of *P. shigelloides*

One hundred and fifty eight suspected *P. shigelloides* were selected for affirmation via molecular method. Only 58 (36.7%) were affirmed as *P. shigelloides* across the two markets as follows: Okitipupa = 30, Igbokoda = 28 (Table 1). They occurred in various percentages in fish storage water (27.6%) fish (37.9%) and Towel (34.5%). *Plesiomonas shigelloides* amplified using the 23S rRNA via Polymerase Chain Reaction is shown in Figure 1. The sequenced *Plesiomonas* species is stated in Table 2.

**Figure I:**
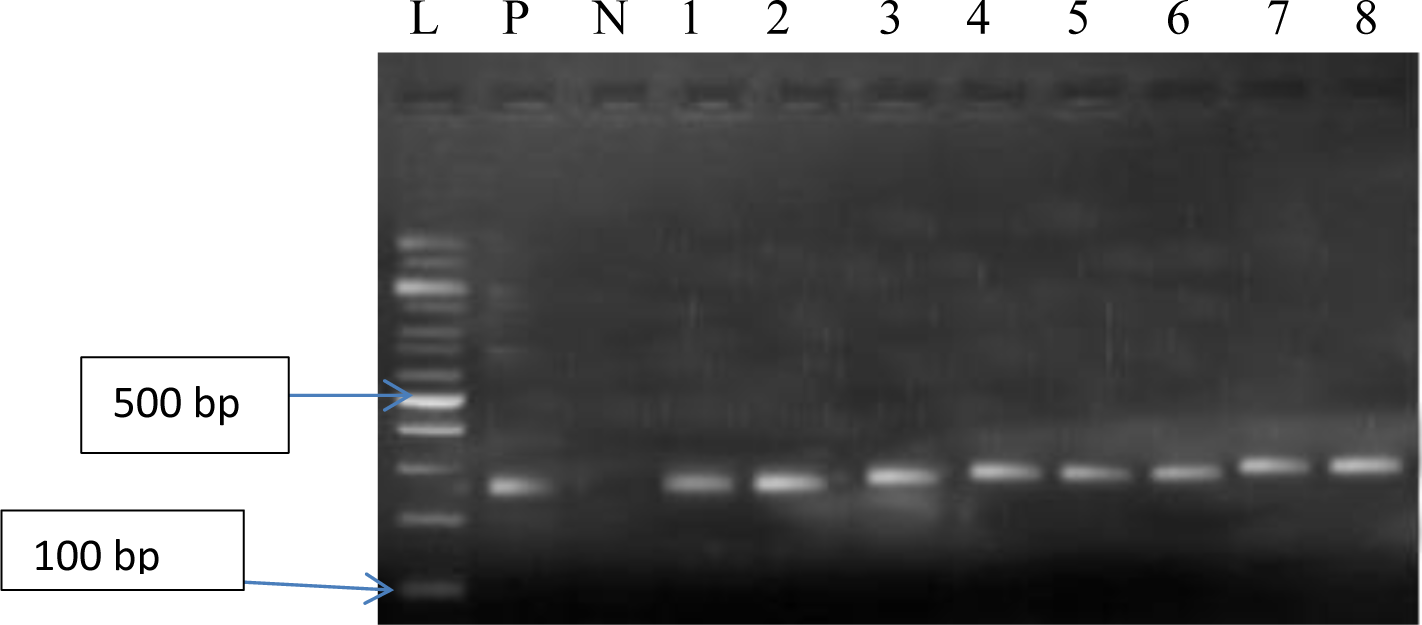
Gel of 284 bp region of *P. shigelloides* (L=100 bp ladder, P = positive control, N = Negative control, 1 – 8 = Positive isolates)

**Table 1:** Distribution of *Plesiomonas shigelloides* across sample sites.

|  | Okitipupa (%) | Igbokoda (%) | Total (%) |
| --- | --- | --- | --- |
| Fish Storage Water (n= 45) | 6 (20) | 10 (35.7) | 16 (27.6) |
| Fish (n= 63) | 12 (40) | 10 (35.7) | 22 (37.9) |
| Towel (n= 50) | 12 (40) | 8 (28.6) | 20 (34.5) |
| Storage water Source (n= 0) | 0 (0) | 0 (0) | 0 (0.0) |
| Total (n =158) | 30 (51.7) | 28 (48.3) | 58 (36.7) |

**Table 2:** Identity of the sequenced *P*. *shigelloides.*

| Isolate | Query cover | % identity | Accession No | Identity |
| --- | --- | --- | --- | --- |
| P5 | 96 | 89.54 | LT575468.1 | *Plesiomonas shigelloides* strain NCTC10360 |
| P13 | 89 | 90.23 | CP027852.1 | *Plesiomonas shigelloides* strain MS-17-188 |
| P27 | 94 | 90.63 | CP050969.1 | *Plesiomonas shigelloides* strain FDAARGOS_725 |
| P36 | 98 | 96.32 | CP062196.1 | *Plesiomonas shigelloides* strain P5462 |
| P45 | 85 | 77.84 | CP087712.1 | *Plesiomonas shigelloides* strain 7A |
| P56 | 92 | 87.57 | LT575468.1 | *Plesiomonas shigelloides* strain NCTC10360 |

### Antibiotics Susceptibility of *P. shigelloides*

All the confirmed *P. shigelloides* were subjected to antibiotic susceptibility test and the study revealed that all the isolates resisted the effect of two or more antibiotics. The antibiotic susceptibility profile of *P. shigelloides* obtained from fish storage water, fish and towel of fish sellers from the two sampled sites are shown in Figure 2 and 3. It was discovered that all the isolates followed the same trend in the two sampled sites. All the isolates obtained from both sites showed resistance to the tested antibiotics in the following trend: Amikacin (21%), Cotrimoxazole (22%), Gentamicin (26%), Ciprofloxacin and Chloramphenicol (34%), Trimethoprim (41%), Streptomycin and Amoxicillin (43%), Meropenem and Vancomycin (45%), Tetracycline (67%), Cefuroxime (71%), Ceftazidime (81%), Ampicillin (83%), Erythromycin (85%). Meanwhile, *P. shigelloides* obtained showed highest susceptibility to Amikacin (79%) followed by cotrimoxazole (78%) and lowest susceptibility to Erythromycin (15%) followed by Ampicillin (17%) (Table 3).

**Figure II:**
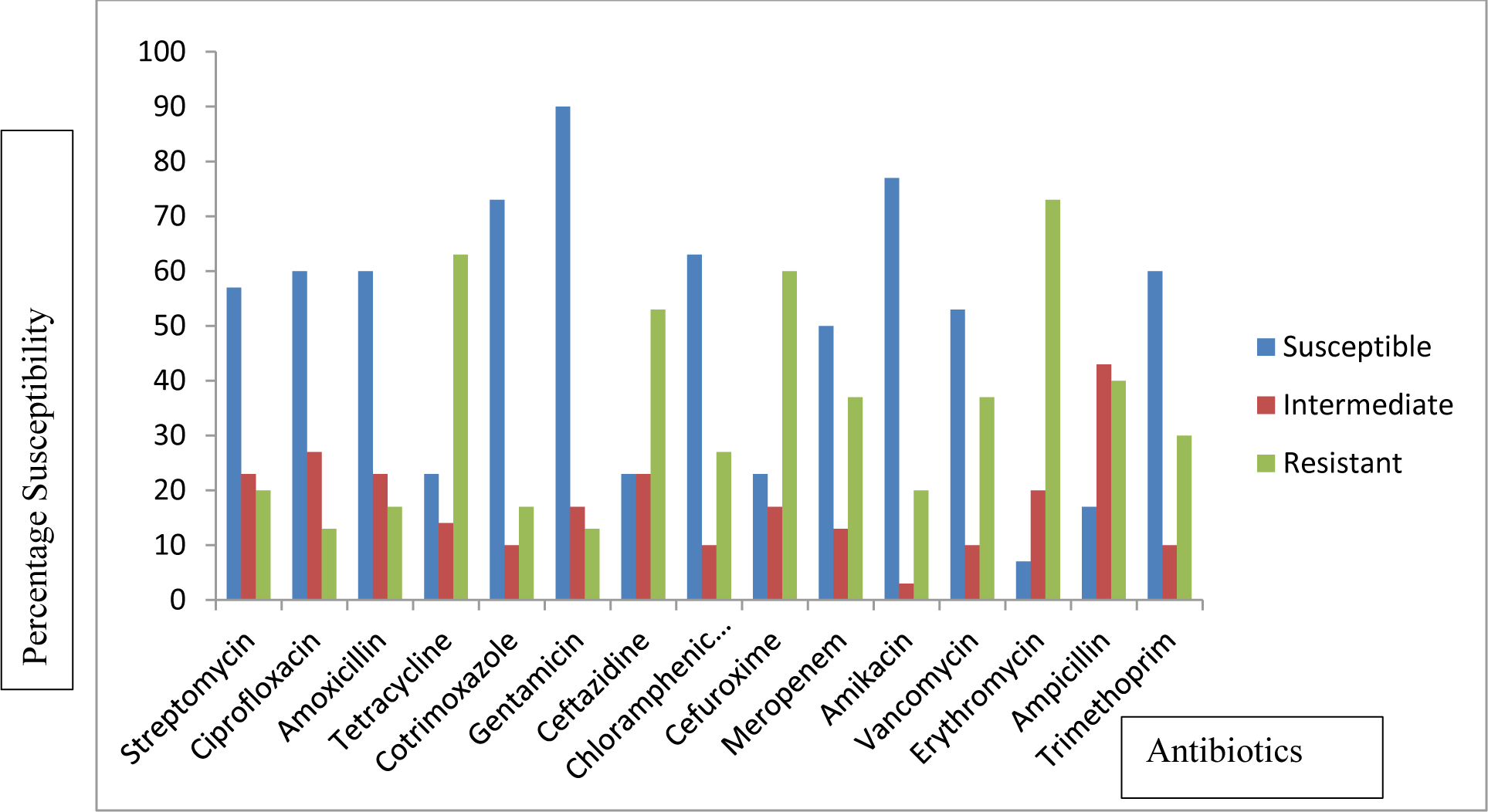
Antibiotics susceptibility profile of *P. shigelloides* isolated from Okitipupa

**Figure III:**
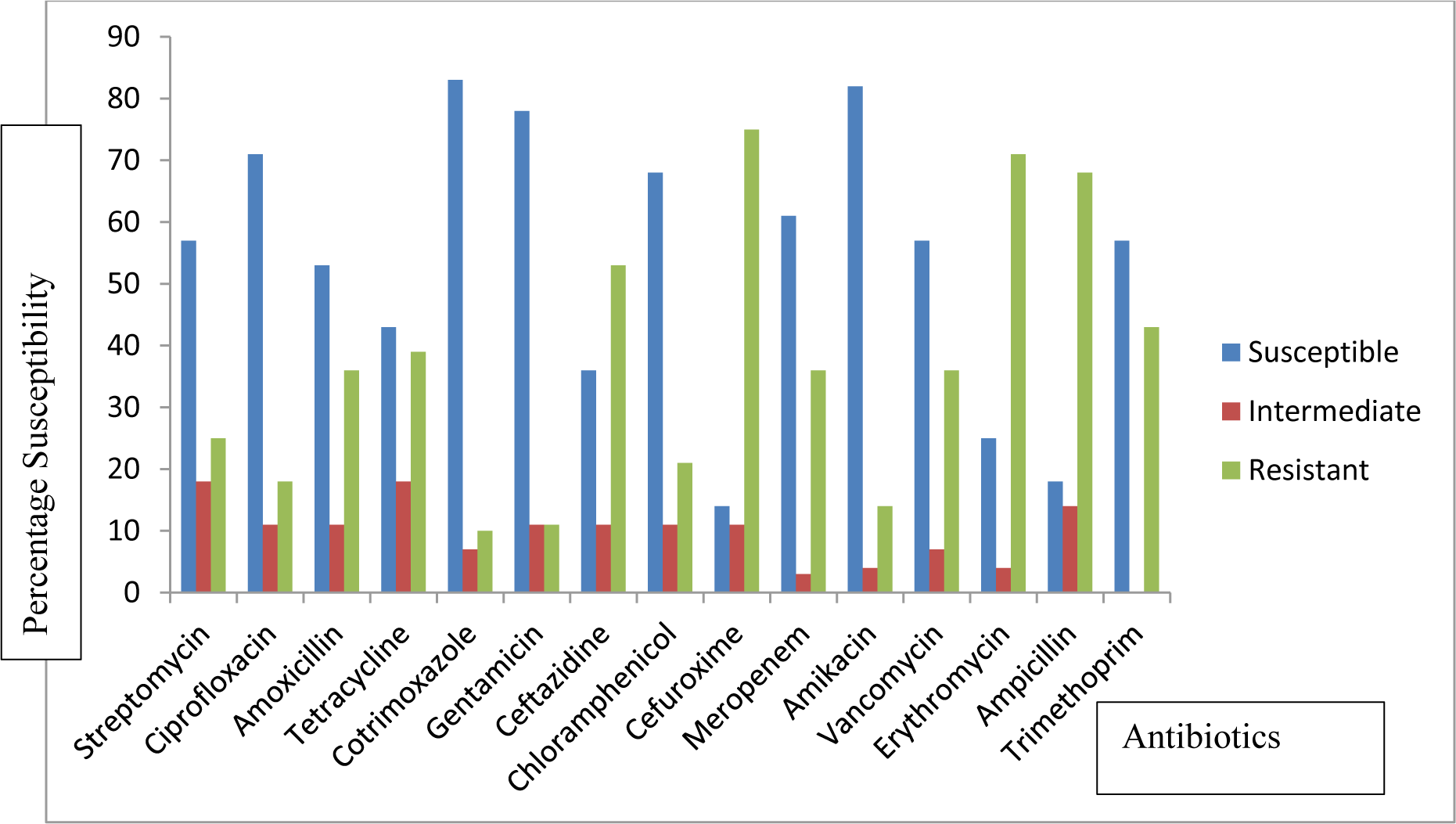
Antibiotics susceptibility profile of *P. shigelloides* isolated from Igbokoda

**Table 3:** Antibiotic resistant profile of *P. shigelloides* isolated from both sites.

|  | STR | CIP | AMO | TET | COT | GEN | CRX | CHL | CPZ | MEM | AMK | VAN | E | AMP | T | Total | MARI |
| --- | --- | --- | --- | --- | --- | --- | --- | --- | --- | --- | --- | --- | --- | --- | --- | --- | --- |
| Okitipupa | 13 | 12 | 12 | 23 | 8 | 9 | 23 | 11 | 23 | 15 | 7 | 14 | 28 | 25 | 12 | 235 | 1.0 |
| Igbokoda | 12 | 8 | 13 | 16 | 5 | 6 | 18 | 9 | 24 | 11 | 5 | 12 | 21 | 23 | 12 | 195 | 1.0 |
| Total | 25 | 20 | 25 | 39 | 13 | 15 | 41 | 20 | 47 | 26 | 12 | 26 | 49 | 48 | 24 | 430 |  |
| % Resistant | 43 | 34 | 43 | 67 | 22 | 26 | 71 | 34 | 81 | 45 | 21 | 45 | 85 | 83 | 41 |  |  |

**Key:** STR- Streptomycin, CIP – Ciprofloxacin, AMO – Amoxicillin, TET – Tetracycline, COT – Cotrimoxazole, GEN - Gentamicin,
, CRX - Cefuroxime, CHL- Chloramphenicol, CPZ - Ceftazidime, MEM- Meropenem, AMK- Amikacin, VAN- Vancomycin,
Trimethoprim, E- Erythromycin AMP- Ampicillin, T- Trimethoprim

### Multiple antibiotic resistance phenotypes (MARPs) of *P. shigelloides*

All *P. shigelloides* isolates resisted more than two (2) antibiotics based on the MARP. Multiple Antibiotics Resistant Phenotypes exhibited by the isolates ranges from resistance to three (3) classes of antibiotics to eight (8) classes of antibiotics with only one isolate obtained from Okitipupa resisting 3 classes of antibiotics. Some of the *P. shigelloides* isolated from fish showed similar antibiotics susceptibility profile with some isolated from fish seller’s towel. The frequent at whom MARPs occurred was high because isolates were resistant to multiple antibiotics ranging from 5 – 11 antibiotics. In total, the two sampling locations’ MAR indices were higher than the threshold value of 0.2, indicating contamination from high-risk sources like overuse of antibiotics in veterinary, human, and agricultural settings, which can unintentionally endanger public health. The computed MARindex was discovered to range from 0.27 to 0.78, with 0.78 being the highest and 0.27 being the lowest, both of which were above the 0.2 limitations (Table 4). This shows that there is a heavy use of antibiotics in the environment and in aquaculture settings.

**Table 4:**
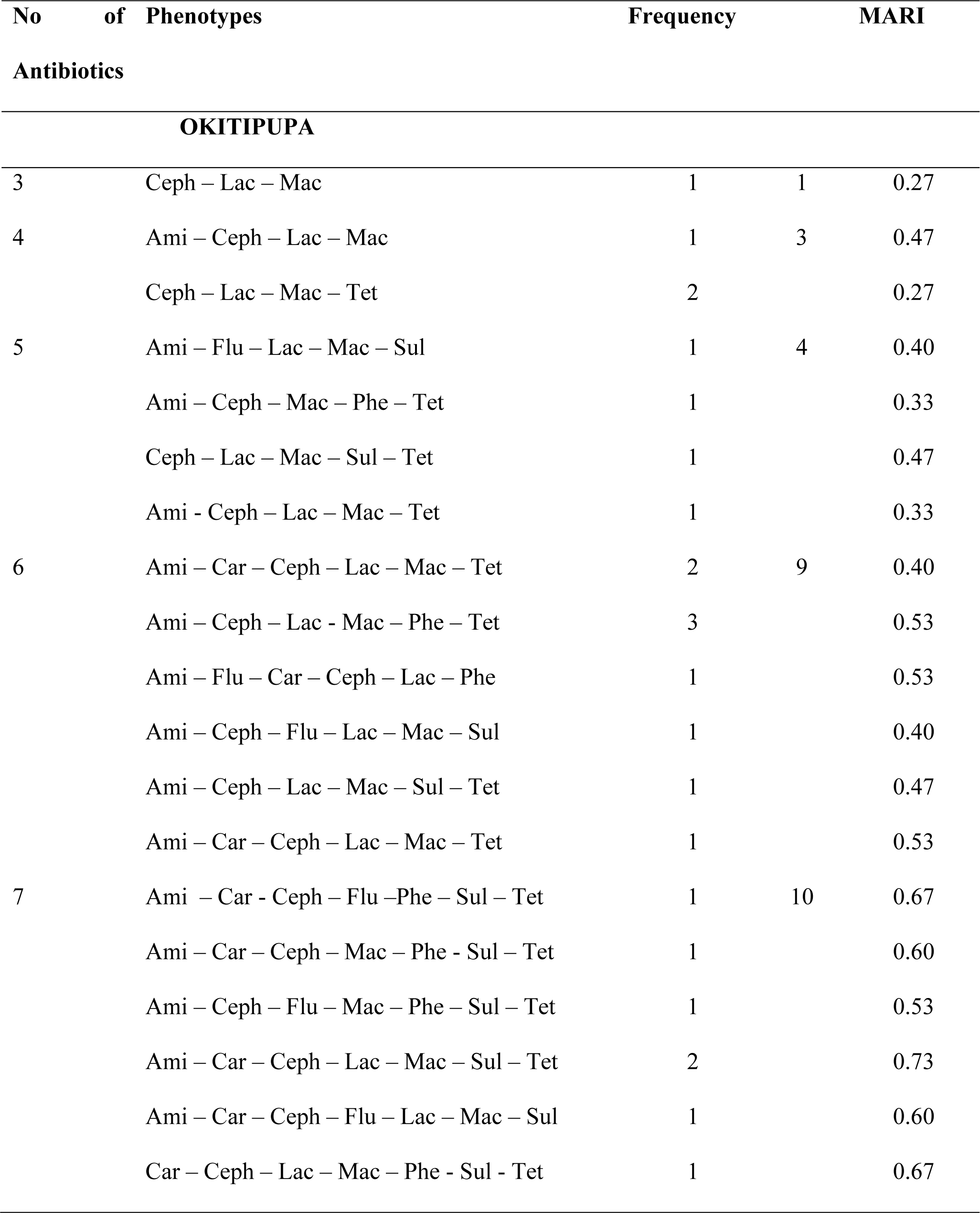
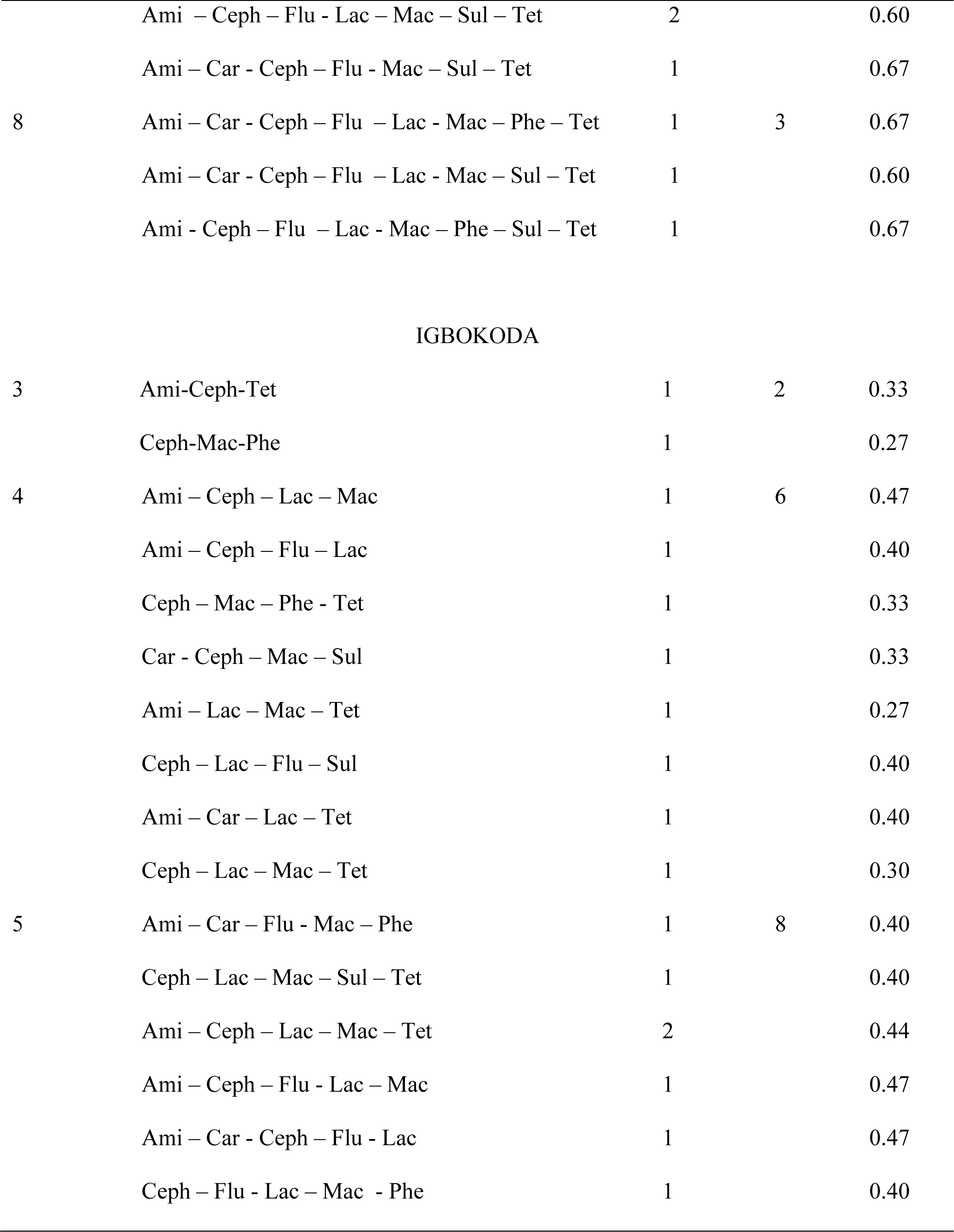
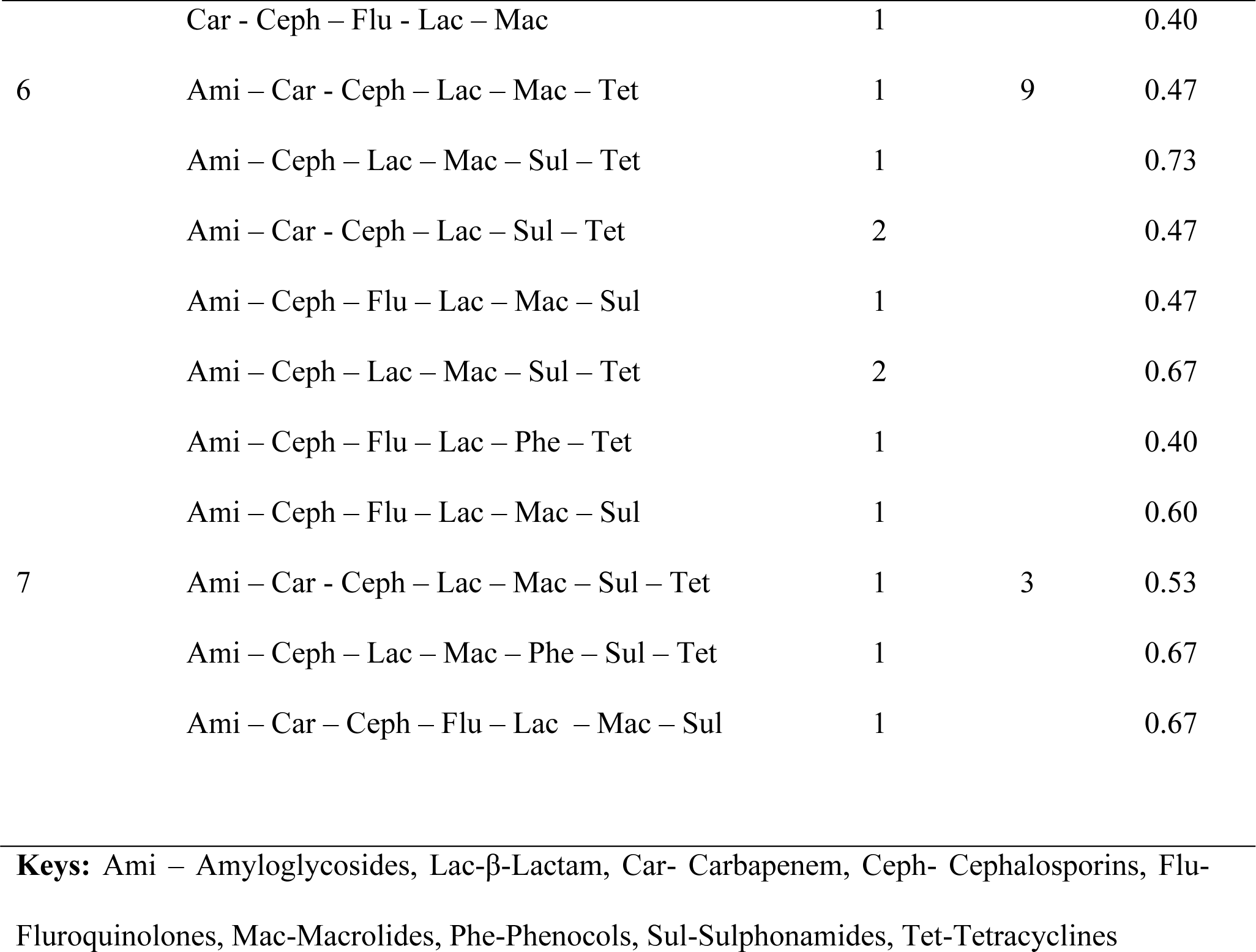
Multiple Antibiotic Resistant Phenotypes exhibited by *P. shigelloides.*

## DISCUSSIONS

Many virulence components, such as adhesiveness, hemolysins, cytotoxins, cholerae-like toxins, and other exoenzymes linked to pathogenicity and are thought to contribute to *Plesiomonas shigelloides* ability to withstand a variety of environmental stresses which could be the reason for their ability to survive in towel (Wang *et al*., 2020). This aquatic bacterium was recognized as the causative agent of a number of opportunistic infectious diseases in both humans and animals that were connected to the water environment. This study examined the antibiogram feature of *P. shigelloides* isolated from fish storage water, fish and fish seller’s towel in Southern part of Ondo State, Nigeria. This was with the view of assessing the ease at which this pathogen can be transmitted from one point to another. Using a species-specific PCR technique, presumptive *P. shigelloides* isolated was validated by amplifying the 23S rRNA gene region between nucleotides C-906 and G-1189 (Adesiyan *et al*., 2019). The presence of such pathogen in fish storage water and fish could be as a result of contaminated source of water used in the aquaculture systems which will definitely affect the quality of fishes reared in the water and this corroborate the findings of Sugital *et al*. (1993) who also isolated *P. shigelloides* in fish intestine as well as the surrounding water. Joh *et al*. (2013) also isolated *P. shigelloides* from aquacultured eels (*Anguilla japonica*) and the surrounding waters in Korean eel farms. Furthermore, Wang *et al*. (2020) also reported *P. shigelloides* as a potential pathogen associated with enteritis in channel catfish which could lead to economic loss for the aquaculture farmers. *P. shigelloides* was found in freshwater bodies, and Adesiyan *et al*. (2019) hypothesized that this might be because of anthropogenic activity near water, effluents being released into rivers, or being washed into them during run-off. Aldova *et al*. (1999) also isolated *P. shigelloides* from water, fish skin swab as well as intestine indicating that this potential pathogen could be termed as one of the normal flora of aquatic environment. These scientists also discovered that a single host harbored several serovars. This study showed that the presence of *P. shigelloides* in the fish samples purchased from both sites could be an indication that the water source is not of good microbiological quality. This study also proof that it is very easy for this pathogen to be transmitted due to its presence in the fish seller’s towel swab. This incidence is a threat to the public health as buyers could use the same towel to clean their hands after purchasing fish and the hands could serve as a transmission vehicle for the pathogen to ready to eat foods in the market that could result in epidemics. The presence of this pathogen could also result in economic loss for farmers due to high mortality rate that could occur in aquatic animals as documented by several researchers (Sugita *et al*., 1993; Rajagopalan *et al*., 2014; Zhang *et al*., 2019; Liu *et al*., 2015; Wang *et al*., 2020; Fuentes-Valencia *et al.,* 2022). When effluent from this aquaculture setting and the waste water from fish sellers in the market are discharged into the environment, it will further contaminate the nearby water bodies via surface run-off hence, a public health threat.

Concern has been expressed by scientists about the emergence of antibiotic resistance in *P. shigelloides* (Adesiyan *et al*., 2019; Ekundayo *et al*., 2020; Wang *et al*., 2020). In this study, it was discovered that all the isolates obtained from both markets followed almost the same trend (Figure 1 and 2) in their response to antibiotics which could be due to the fact that the same type of fish were assessed or the same source of fish being sold in the market at the time of sampling or closeness of the area of the aquaculture system where fishes was been purchased that might have exposed them to the same environmental conditions. It was discovered that *P. shigelloides* isolates revealed varying levels of resistance to several antibiotics (Table 3). The isolates showed varying levels of resistance to erythromycin, tetracycline, ampicillin, chloramphenicol, amoxicillin, and streptomycin, which was consistent with other researchers’ observations (Abdelhamed *et al*., 2018; Wang *et al*., 2020). This study also corroborate the findings of Fuentes-Valencia *et al.,* (2022) who reported resistance to several antibiotics including carbapenem and cephalosporin in *P. shigelloides* they isolated from *Oncorhynchus chrysogaster*. Resistance observed in this study also agrees with the findings of Adesiyan *et al*., (2019) who recorded resistance in *P. shigelloides* isolated from selected surface water in southwest Nigeria but disagree with the susceptibility recorded against meropenem because 45 % of the *P. shigelloides* isolated in this study resisted the effect of meropemem and this could be attributed to diversity in the source of isolation as well as geographical location. Resistance in bacteria may be acquired via horizontal gene transfer from other bacterial species, animal or human origin contaminating the water bodies used in the farm (Fuentes-Valencia *et al.,* 2022). Additionally, it was suggested that the *Plesiomonas* resistance might be due to the organism’s outer membrane blocking the entry of antibiotics into its cell wall, suggesting that this organism may be naturally resistant to various antibiotics, particularly penicillin. Some of the isolates obtained from towels of fish sellers showed antibiotics susceptibility profile similar to those isolated from fish which demonstrated the ease at which this emerging pathogen can be transmitted from one place to another hence constitute a public health threat especially in immuno-compromised individual. In contrast, isolates varied in their susceptibility to gentamicin and ciprofloxacin, which is consistent with earlier research (Kim *et al*., 2015; Adesiyan *et al*., 2019). The susceptibility of this organism to aminoglycosides may vary depending on its sources, as evidenced by reports that clinical isolates of this organism are less gentamicin susceptible than environmental isolates of this species (Chen *et al*., 2013; Nwokocha and Onyemelukwe, 2014; Pence, 2016; Lee *et al*., 2016). According to reports of human gastroenteritis caused by *P. shigelloides* in immunocompetent and immunocompromised individuals, the resistance to some routinely used antibiotics found in this study should be a major public health concern. Based on the multiple antibiotics resistance phenotypes exhibited by isolates obtained from both sample sites, it was discovered that isolates obtained from Igbokoda resisted the effect of antibiotics ranging from 3 – 7 classes of antibiotics. Meanwhile, those obtained from Okitipupa ranged from 3 – 8 classes of antibiotics and three isolates were found to show resistant to 8 classes of antibiotics. One of the isolates that were resistant to 8 classes of antibiotics was gotten from fish sample while the remaining two (2) were obtained from the towel and this could be as a result of diversity in the strain. Variation in the antibiotics susceptibility profile exhibited by isolates obtained in this study could be as a result of strain diversity as previously mentioned by other researcher who obtained several serovars of *P. shigelloides* in a single host (Aldova *et al*., 1999).

The two sampled locations had *P. shigelloides* isolates with high MARindex values that above the established threshold value of 0.2 for identifying low- and high-risk contamination (Krumperman, 1983). Both the contaminated water source and the overuse of antibiotics in aquaculture facilities may be to blame for the high MARindex figures observed in the two sampled marketplaces. Numerous studies have shown the part played by wastewater effluent in the spread of enteric bacteria containing various antibiotic-resistant plasmids (Silva *et al*., 2006; Akhter *et al*., 2014) because of the amount of nutrients and high concentration of microbes, and a significant location for horizontal gene transfer (Sloan *et al*., 2014). Because of the frequent and unauthorized use of antibiotics, the isolates represent high risk sources of contamination, as indicated by the high MAR score.

## CONCLUSION

The presence of *P. shigelloides* in the fish storage water and fish samples may be an indication for future outbreak of gastrointestinal infection via consumption of contaminated fish. However, chances of cross contamination of other food products in the market with fish seller’s towel serving as transmission vehicle as well as through kitchen materials could result in infectious diseases of great public health concern. The high incidence of resistant *P. shigelloides* species against amyloglycosides, cephalosporins, flouroquinolone, macrolide, tetracycline and β-lactam implies increased resistance against some antibiotics of choice which could be a threat to the effective treatment of infection associated with *P. shigelloides* in the study area. This suggests abuse of antibiotics usage around the study areas and aquaculture settings. Therefore, this study showed that the fish sold in this area serve as potential reservoirs of multidrug resistant *P. shigelloides* that may cause outbreaks of diarrhea associated infection in Southern part of Ondo State, Nigeria and this opportunistic pathogen can easily be transmitted via inanimate object. Hence, we recommend proper processing of fresh fish and the practice of good hygiene for both sellers and buyers so as to prevent infectious diseases that may be associated with the pathogen.

## IMPACT STATEMENT

This research work showed that the pathogen in catfish commonly sold in the market can be transmitted via the fish seller’s towel and transferred to human through direct contact and cross contaminate other ready to eat foods thereby resulting in a potential public health challenge.

## DATA AVAILABILITY

All data relating to this manuscript are available in the manuscript

## ETHICAL APPROVAL

This study neither utilized human study nor animal’s model. Hence, does not require any ethical approval.

## AUTHOR CONTRIBUTIONS

Temitope D. Agboola: Conceptualized, Visualize, Supervised and draft the Manuscript. Monica O. Oguntimehin: Involved in Investigation, Data Curation and review of the the manuscript. Oluwapelumi R. Oyeneye and Isaac O. Obadofin: Involved in investigation, data curation and formal analysis. All authors agreed on the final manuscript.

## ACKNOWLEDGEMENTS

The authors appreciate the market women in the two communities for their cooperation during sample collection and the head, Department of Biological Sciences, Olusegun Agagu University of Science and Technology Okitipupa, Ondo State Nigeria for providing enabling environment to carry out this research. We are also grateful to the Director and staff of Central Science Laboratory, Obafemi Awolowo University Ile – Ife for allowing us to carry-out some aspect of this study.

